# Prognostic value of Red Blood Cell Distribution Width in Acute Kidney Injury Patients: A Systematic Review and Meta-analysis

**DOI:** 10.1101/750190

**Authors:** Zhengsheng Liu, Shanshan Wang, Xiongbo Yao, Jinchun Xing

**Author notes:** Corresponding author: Department of Urology, The First Affiliated Hospital of Xiamen University, Siming District, Xiamen, Fujian, China. Electronic address.

## Abstract

**Background:** The conflicting result with regard to Red Cell Distribution Width (RDW) with Acute Kidney Injury (AKI) has been reported. This systematic review and meta-analysis were aimed to investigate RDW and prognostic value in AKI patients.

**Methods/Main Results:** This meta-analysis included 1251 cases and 1663 controls with a total of 7 enrolled published papers. The results of RDW levels were significantly associated with patients of AKI (WMD=1.127, 95% CI=0.426-1.827; P=0.002), with statistically significant heterogeneity (I^2^ =95.80%, P_heterogeneity_ =0.000, random-effects model).

**Conclusions:** In conclusion, the results of this present meta-analysis suggest that the RDW value is a positive prognostic indicator in patients with AKI. However, these results were obtained on the basis of RCS or small sample sizes studies. Further functional studies with additional data would be needed to validate our findings.

## Introduction

Acute kidney injury (AKI), one of the most common and serious diseases in the intensive care unit (ICU), is defined as a drastic decrease in glomerular filtration rate with increase in serum creatinine (SCr) or decrease in urine output[1]. Previous studies have confirmed that AKI is an independent risk factor for high mortality range from 36.4% to 50%[1–4], especially in patients with critically illness[5]. In addition, patients with renal function recovery were often suffered from the chronic kidney disease and the major cardiovascular adverse events [6, 7]. Therefore, the early identification of the patients who may progress to AKI and use preventive measures to prevent the occurrence of AKI is crucial. However, AKI was only monitored by SCr which is elevated several hours to days after the renal function loss more than 50%[8, 9].

The red cell distribution width (RDW) is a quantitative measure of variability in the size of circulating erythrocytes with higher values reflecting greater heterogeneity in cellular sizes. As a part of complete blood cell count, RDW is generally determined among hospitalized patients because of its simplicity and availability[10, 11]. RDW is typically used to differentiate the different causes of anemia[12], however, the present studies have observed that RDW could be used as a predictive marker in various diseases, including cardiovascular disease [11, 13], malignant tumors[14, 15], acute pancreatitis[16].

Currently, RDW was reported to be an independent predictor in patients with AKI [17–20] and the result remains inconclusion. Our meta-analysis was conducted to screen the current literature to evaluate prognostic value of RDW in the patients with AKI and explain this conflicting result.

## Methods

### Search strategy

This systematic review was based on the Preferred Reporting Items for Systematic Reviews and Meta-Analyses (PRISMA) statement guidelines. The association between RDW and AKI was evaluated by two investigators independently. The following databases were electronically searched on 12/6/2019, including PubMed, Embase (host: OVID) from 1980 to June 2019, and Web of Science. The following keywords of the search strategy were used: “acute kidney injury” or “acute kidney failure” or “acute renal injury” or AKI and “red cell distribution width” or RDW.

### Inclusion and exclusion criteria

For those studies which were strictly qualified the following criteria were included: (a) studies focus on the association between RDW and AKI; (b) reported the available data on Means and SD values or interquartile range for RDW levels in AKI patients; (c) The diagnosis of AKI was clear; and (d) provided clear clinically relevant outcomes according to the comparisons of AKI versus non-AKI. The studies that met exclusion criteria was excluded and the following was exclusion criteria: (a) not research articles, for example conference abstracts, books, posters, letters, and review articles; (b) studies on animals not human; and(c) underlying disease with early renal injury.

### Data Extraction

Based on the previous inclusion and exclusion criteria, the detail information and data of each eligible study was extracted by two review authors (Liu and Wang). An experienced author (Yao) participated to resolve controversy if the outcome and conclusion of the study still remained some uncertainties or discrepancies.

In this meta-analysis, data and information of the individuals of included studies were extracted, including the first authors name, publication year, country, ethnicity (Caucasian, Asian), study design(retrospective cohort study and case-control study), the mean age of the case and control (years), study population, the total (male and female)and male numbers of cases and controls, definition of AKI, the RDW levels (means and standard deviation (SD) or median and interquartile range) of cases and controls, definition of outcomes, the quality of each study.

### Quality Assessment

According to the guidelines of the NewCastle-Ottawa Quality Assessment Scale (NOS, http://www.ohri.ca/programs/clinical_epidemiology/oxford.asp), the quality of the eligible study was assessed. The NOS Cohort Studies for retrospective cohort study (RCS) score system was employed by corresponding to various type of study. Same to date extraction, two review authors did this work first, then the third author resolved controversy when the study still remained some uncertainties. The total NOS quality scores ranged from 0-9, and the NOS scores higher than 5 points indicated better quality and were suitable for meta-analysis.

### Statistical analysis

We evaluated the association between RDW and AKI by extracting the mean and standard of RDW levels in each study. The published article of Van Driest et al. [21], which include two studies, did not provide the precise mean and SD values. We attempted to correlate this corresponding author but failed. Therefore, means and SD values in this study was calculated on the basis of the median and interquartile range.

By using random-effects model (DerSimonian and Laird method) or the fixed-effects model (Mantel and Haenszel method), the weighted mean difference (WMD) and 95% CIs were obtained to evaluate the prognostic value of RDW in AKI patients. All statistical analysis was performed using Stata 12.0 software (StataCorp, College Station, Texas, USA). The statistical significance was finally defined as a 2-tailed P<0.05.

To obtain the pooled results of heterogeneity in each study, we performed Cochran’s Q test and I-squared statistic[22]. For heterogeneity, P<0.10 in Q test and/or I^2^ value>50% was considered to indicate statistical significance, and the random-effects model (DerSimonian and Laird method) was applied to estimate the summary WMD and 95% CI; otherwise the fixed-effects model (Mantel and Haenszel method) was applied[23]. The sensitivity analysis was also carried out to evaluate the stability of our meta-analysis by omitting one study in total studies. Additionally, subgroup analyses were stratified by ethnicity(Caucasian, Asian), baseline mean age (years) of AKI(≥ 60, <60), Study population(CI-AKI, CSA-AKI, Other), Sample size in AKI cases(≥ 100, <100). Finally, we performed the Begg’s [24] funnel plot and Egger’s[25] tests to explore the potential publication bias, and the P value < 0.05 was considered statistically significant.

## Results

### Study characteristics

A flow diagram of the data search and study selection is presented in Figure 1. We carefully screened the databases and found a total of 96 articles (PubMed= 37, Embase= 32, Web of Science= 27) in initial search process. With 32 duplicate articles, which were removed, and leaving 64 articles for examining and reading titles and abstracts. Then, the full texts 22 papers were downloaded and examined for further investigation. Finally, a total of 7 enrolled published papers (8 studies) included 1251 cases and 1663 controls were identified meeting the criteria for inclusion in our meta-analysis, with publication years ranging from 2015 to 2018[21, 26–31].

**Figure 1:**
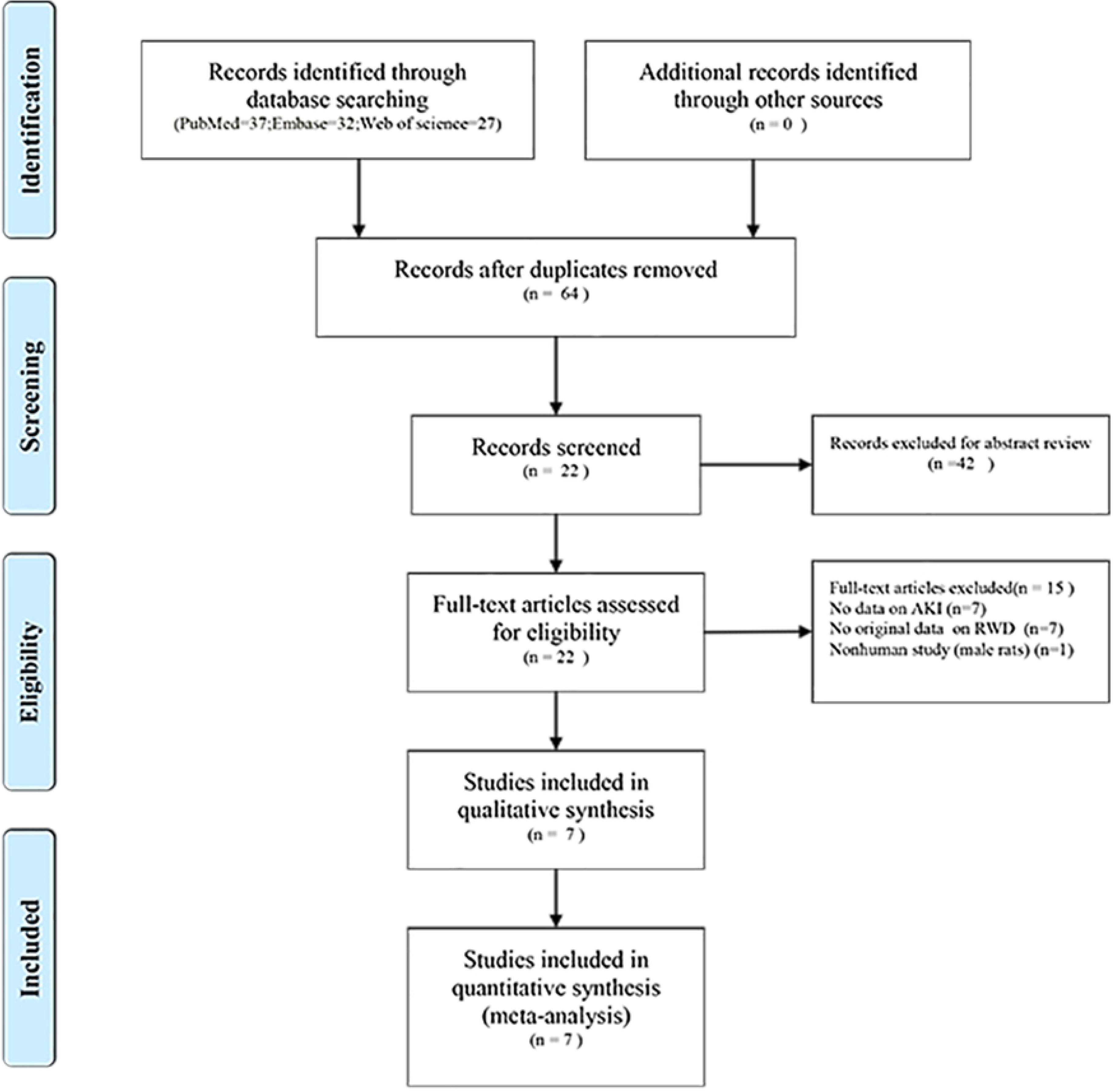
Flow chart of study selection; RDW, AKI, acute kidney injury; red blood cell distribution width.

The key features of 8 studies on the association between RDW and AKI are shown in Table 1. In those studies, half of which including Caucasian descents (3 published papers included 4 RCSs)[21, 26, 27] and the other half of which including Asian descents[28–31]. All the included studies were prospective cohort studies, and the mean age years of the patient with AKI range from 6.7 to 68.9. For the study population, 3 focused on contrast-induced acute kidney injury (CI-AKI), 3 focused on cardiac surgery-associated acute kidney injury (CSA-AKI), the other 2 focused on children with sepsis and patients who were receiving extracorporeal membrane oxygenation. The quality assessment of NOS scores range from 7 to 9, which demonstrated that it was suitable for meta-analysis.

### Main results and Sub-group analyses

All the availability data from the included RCS were synthesized to evaluate the association between the RDW levels and AKI. The results of RDW levels were significantly associated with patients with AKI (WMD=1.127, 95% CI=0.426-1.827; P=0.002), with statistically significant heterogeneity (I^2^ =95.80%, Pheterogeneity =0.000, random-effects model; Figure. 2). Then, sub-group analyses were performed according to ethnicity (Caucasian, Asian), baseline mean age years of AKI(≥ 60, <60), Study population(CI-AKI, CSA-AKI, Other), Sample size in AKI cases(≥ 100, <100), and the detailed information was shown in Table 2. When stratifying according to ethnicity, the statistically significant associations between RDW levels and patients with AKI were observed both in Asians (WMD= 1.472, 95% CI=0.392-2.552; P=0.008) and Caucasian (WMD= 0.577, 95% CI=0.308-0.846; P=0.000). For mean age years, a significant association was observed in both AKI patients baseline mean age over 60 years (WMD= 0.725, 95% CI=0.216-1.233; P=0.005) and under 60 years (WMD= 1.941, 95% CI=0.226-3.656; P=0.027). In the study population, the results showed that the RDW levels were of significant prognostic value with AKI patients, including CI-AKI (WMD= 0.582, 95% CI=0.314-0.849; P=0.000), CSA-AKI (WMD= 1.297, 95% CI=1.174-1.420; P=0.000), except the group of others (WMD= 1.751, 95% CI=-0.895-4.397; P=0.195; Table 2). The WMD of RDW levels was significant in studies with sample size more than 100 (WMD= 0.912, 95% CI=0.202-1.623; P=0.012), however, no significant prognostic value was detected in studies with sample size less than 100 (WMD= 1.261, 95% CI=-0.320-2.842; P=0.118).

**Figure 2.**
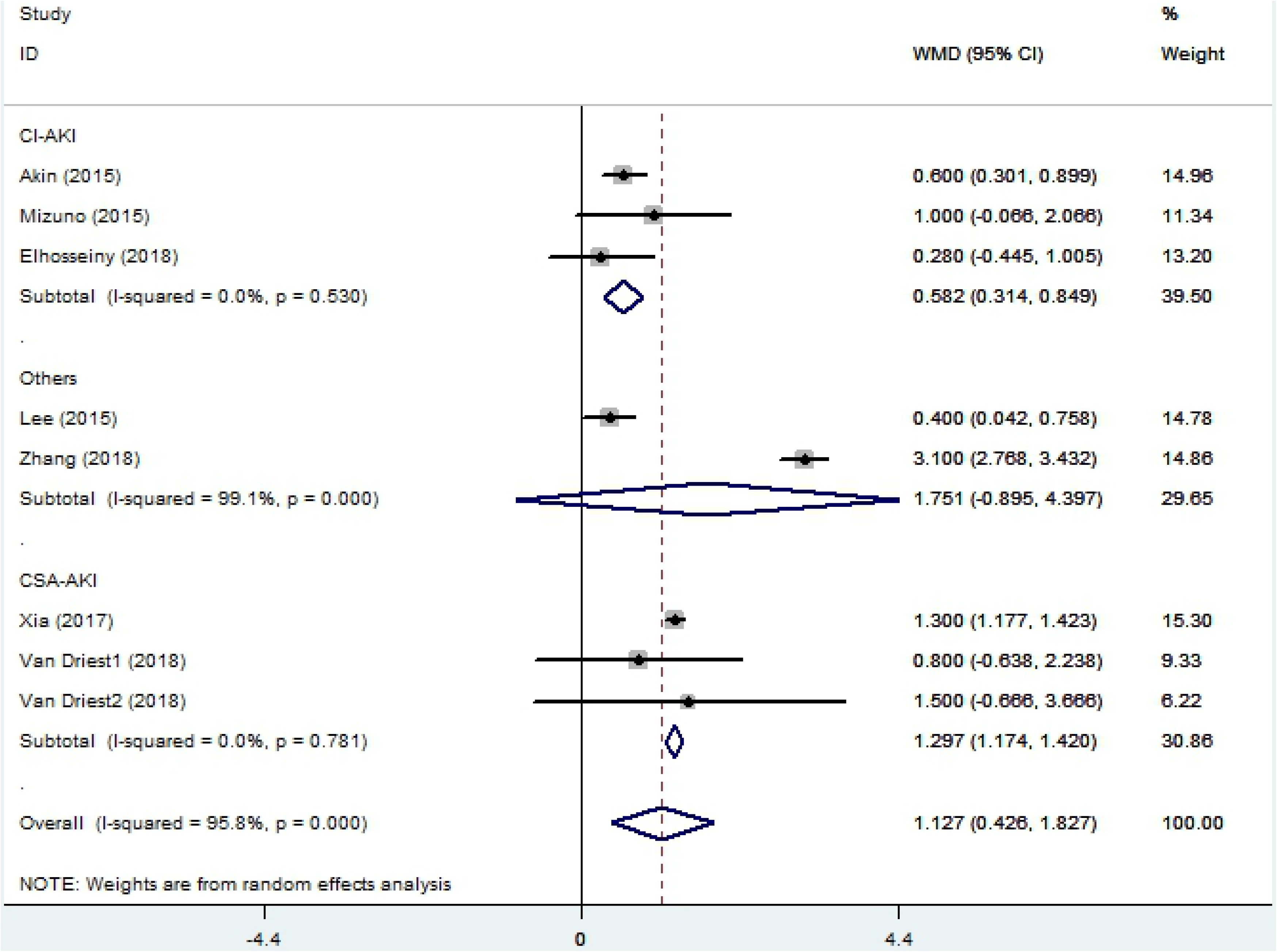
Forest polt

### Heterogeneity and Sensitivity analyses

As was shown in Table 2, we detected significant heterogeneity in the meta-analysis, however, there was no statistically significant heterogeneity according to Caucasian (I^2^= 0.00%; P_heterogeneity_=0.692); CI-AKI (I^2^= 0.00%; P_heterogeneity_=0.530); CSA-AKI (I^2^= 0.00%; P_heterogeneity_=0.781). Then we performed sensitivity analyses by consistently omitting one study to evaluate the influence of each single study on results, and it indicated that the results of this metaanalysis was reliable (Figure. 3). Furthermore, Begg’s funnel plot and Egger’s test were used to evaluate the publication bias. And there was no publication bias detected in association with RDW and AKI patients, with Begg’s funnel plot (P_Begg_= 0. 902) and Egger’s regression test (P_Egger_ = 0.824).

**Figure 3.**
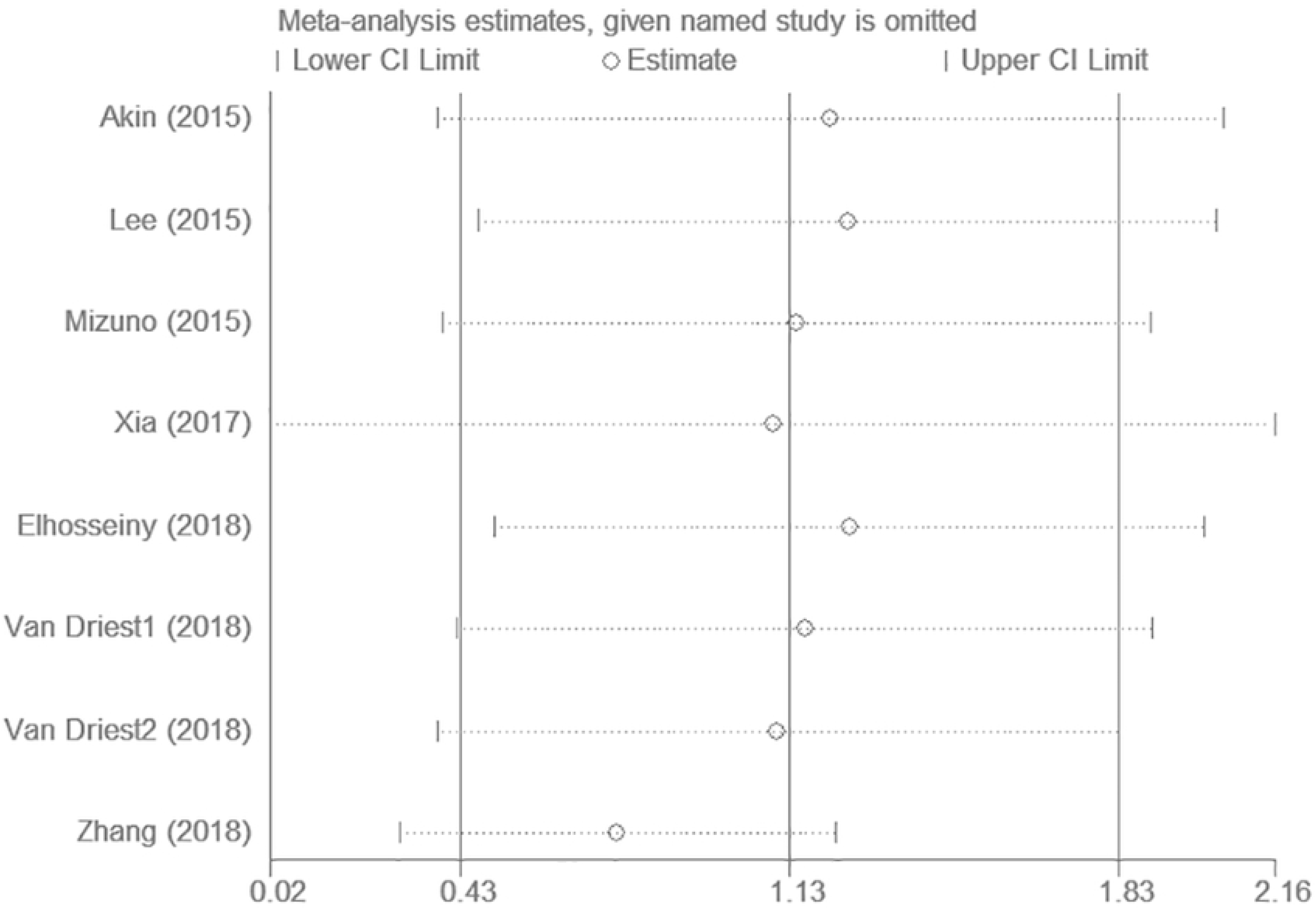
WMD pooled value

## Discussion

AKI is a common complication of open heart surgery[32], major abdominal surgery[33] and contrast-associated diseases[34], with similar risk factor and outcome associations across different surgery type[35]. In the current study, AKI is frequently transient condition and strongly associated with increased mortality and morbidity[36]. As far as we know, AKI is interconnected with Chronic Kidney Disease (CKD), which results in end-stage renal disease risk and even death. The rates of death associated with CSA-AKI is maintained as high as 10 years regardless of other risk factors, even for those patients who had fully recovered from renal injury[37]. Although the most surgical treatment of secondary AKI is mild, it is still associated with long-term risk of death, especially in post-operative patients with severe AKI.

RDW is a rapid, simple, routinely laboratory test as a part of complete blood count, which is used to evaluate the abnormal properties of red blood cells and the possible causes of anemia. To our knowledge, high RDW is concerned with several diseases, including cardiovascular diseases[38, 39]; chronic kidney disease[40]; esophageal cancer[41]; upper aerodigestive tract cancer[42]; breast cancer[43]. Low RDW, however, may not be clinically significant. Recently, the predictive value of red blood cell distribution width for acute kidney injury was performed in several studies[44, 45]. RDW is considered as an independent prognostic marker in AKI patients, more importantly, higher RDW means high risk of mortality in AKI[17, 18]. Although RDW are known to be associated with AKI, the result remains unclear.

In present meta-analysis, the results demonstrated that RDW value are significantly higher in patients with AKI compared with that of non-AKI (WMD=1.127, 95% CI=0.426-1.827; P= 0.002). In subgroup analysis, the significant association was observed in CI-AKI (WMD= 0.582, 95% CI=0.314-0.849; P=0.000), and CSA-AKI (WMD= 1.297, 95% CI=1.174-1.420; P=0.000), with no heterogeneity detected. We also observed significant value in the ethnicity, mean age, and sample size in AKI cases. Collectively, this meta-analysis indicated that RDW would be prognostic for patients with AKI, especially for CI-AKI and CSA-AKI.

Although the mechanism of the RDW values in patients with AKI is at present unclear, several potential explanations may account for the relationship between RDW and AKI. Among the risk factors, anemia is the independent related to AKI[46], one present study indicated that the elevated RDW group in AKI was found to have low red blood cell (RBC) which may contribute to increase AKI occurring[28]. Previous studies have demonstrated RDW levels are associated with inflammation and serum antioxidants which mediate effects on RDW through IL-6[47], and the IL-6 levels in serum may predict the mortality of AKI patients[48]. In addition, the increased of RDW levels was involved in the impaired residual renal functions[49].

Meanwhile, there are some weaknesses and limitations that should be emphasized in current study. First, we cannot disregard that the heterogeneity across the eligible study was observed in ethnicity, mean age, and sample size. Second, the study design of included studies was totally RCS, and the small number of included studies with a limited number of patients was selected for the present meta-analysis. Third, lack of consolidated standards of AKI, and the measurements of RDW is sometime completely discordant in those included studies. Forth, the patients receiving ECMO, which probably have affected on RDW values, could not be exactly evaluated. Finally, due to absence of required information, we did not have an opportunity to understand the mortality, outcome event, presence or absence of anemia.

## Conclusions

In conclusion, the results of this present meta-analysis suggest that the RDW value is a positive prognostic indicator in patients with AKI. However, these results were obtained on the basis of RCS or small sample sizes studies. Further functional studies with additional data would be needed to validate our findings.

## Authors’ Contributions

Zhengsheng Liu and Shanshan Wang contributed equally to this work as first authors.

## References

1. Bellomo R, Kellum JA, Ronco C. Acute kidney injury. Lancet (London, England). 2012;380(9843):756–66. Epub 2012/05/24. doi: 10.1016/s0140-6736(11)61454-2. PubMed PMID: 22617274.

2. Uchino S, Kellum JA, Bellomo R, Doig GS, Morimatsu H, Morgera S, et al. Acute renal failure in critically ill patients: a multinational, multicenter study. Jama. 2005;294(7):813–8. Epub 2005/08/18. doi: 10.1001/jama.294.7.813. PubMed PMID: 16106006.

3. Joannidis M, Metnitz B, Bauer P, Schusterschitz N, Moreno R, Druml W, et al. Acute kidney injury in critically ill patients classified by AKIN versus RIFLE using the SAPS 3 database. Intensive care medicine. 2009;35(10):1692–702. Epub 2009/06/24. doi: 10.1007/s00134-009-1530-4. PubMed PMID: 19547955.

4. Hoste EA, Bagshaw SM, Bellomo R, Cely CM, Colman R, Cruz DN, et al. Epidemiology of acute kidney injury in critically ill patients: the multinational AKI-EPI study. Intensive care medicine. 2015;41(8):1411–23. Epub 2015/07/15. doi: 10.1007/s00134-015-3934-7. PubMed PMID: 26162677.

5. de Mendonca A, Vincent JL, Suter PM, Moreno R, Dearden NM, Antonelli M, et al. Acute renal failure in the ICU: risk factors and outcome evaluated by the SOFA score. Intensive Care Med. 2000;26(7):915–21. Epub 2000/09/16. PubMed PMID: 10990106.

6. Coca SG, Singanamala S, Parikh CR. Chronic kidney disease after acute kidney injury: a systematic review and meta analysis. Kidney international. 2012;81(5):442–8. Epub 2011/11/25. doi: 10.1038/ki.2011.379. PubMed PMID: 22113526; PubMed Central PMCID: PMCPMC3788581.

7. Schiffl H, Fischer R. Five-year outcomes of severe acute kidney injury requiring renal replacement therapy. Nephrology, dialysis, transplantation: official publication of the European Dialysis and Transplant Association - European Renal Association. 2008;23(7):2235–41. Epub 2008/04/15. doi: 10.1093/ndt/gfn182. PubMed PMID: 18408072.

8. Herget-Rosenthal S, Marggraf G, Husing J, Goring F, Pietruck F, Janssen O, et al. Early detection of acute renal failure by serum cystatin C. Kidney international. 2004;66(3):1115–22. Epub 2004/08/26. doi: 10.1111/j.1523-1755.2004.00861.x. PubMed PMID: 15327406.

9. Haase M, Devarajan P, Haase-Fielitz A, Bellomo R, Cruz DN, Wagener G, et al. The outcome of neutrophil gelatinase-associated lipocalin-positive subclinical acute kidney injury: a multicenter pooled analysis of prospective studies. Journal of the American College of Cardiology. 2011;57(17):1752–61. Epub 2011/04/23. doi: 10.1016/j.jacc.2010.11.051. PubMed PMID: 21511111; PubMed Central PMCID: PMCPMC4866647.

10. Goyal H, Awad H, Hu ZD. Prognostic value of admission red blood cell distribution width in acute pancreatitis: a systematic review. Annals of translational medicine. 2017;5(17):342. Epub 2017/09/25. doi: 10.21037/atm.2017.06.61. PubMed PMID: 28936436; PubMed Central PMCID: PMCPMC5599272.

11. Tonelli M, Sacks F, Arnold M, Moye L, Davis B, Pfeffer M. Relation Between Red Blood Cell Distribution Width and Cardiovascular Event Rate in People With Coronary Disease. Circulation. 2008;117(2):163–8. Epub 2008/01/04. doi: 10.1161/circulationaha.107.727545. PubMed PMID: 18172029.

12. Thompson WG. Red cell distribution width in alcohol abuse and iron deficiency anemia. Jama. 1992;267(8):1070–1. Epub 1992/02/26. doi: 10.1001/jama.1992.03480080040013. PubMed PMID: 1735918.

13. Felker GM, Allen LA, Pocock SJ, Shaw LK, McMurray JJ, Pfeffer MA, et al. Red cell distribution width as a novel prognostic marker in heart failure: data from the CHARM Program and the Duke Databank. Journal of the American College of Cardiology. 2007;50(1):40–7. Epub 2007/07/03. doi: 10.1016/j.jacc.2007.02.067. PubMed PMID: 17601544.

14. Li Z, Hong N, Robertson M, Wang C, Jiang G. Preoperative red cell distribution width and neutrophil-to-lymphocyte ratio predict survival in patients with epithelial ovarian cancer. Scientific reports. 2017;7:43001. Epub 2017/02/23. doi: 10.1038/srep43001. PubMed PMID: 28223716; PubMed Central PMCID: PMCPMC5320446.

15. Montagnana M, Danese E. Red cell distribution width and cancer. Annals of translational medicine. 2016;4(20):399. Epub 2016/11/22. doi: 10.21037/atm.2016.10.50. PubMed PMID: 27867951; PubMed Central PMCID: PMCPMC5107391.

16. Zhang T, Liu H, Wang D, Zong P, Guo C, Wang F, et al. Predicting the Severity of Acute Pancreatitis With Red Cell Distribution Width at Early Admission Stage. Shock (Augusta, Ga). 2018;49(5):551–5. Epub 2017/09/16. doi: 10.1097/shk.0000000000000982. PubMed PMID: 28915220.

17. Oh HJ, Park JT, Kim JK, Yoo DE, Kim SJ, Han SH, et al. Red blood cell distribution width is an independent predictor of mortality in acute kidney injury patients treated with continuous renal replacement therapy. Nephrology, dialysis, transplantation: official publication of the European Dialysis and Transplant Association - European Renal Association. 2012;27(2):589–94. Epub 2011/06/30. doi: 10.1093/ndt/gfr307. PubMed PMID: 21712489.

18. Wang B, Lu H, Gong Y, Ying B, Cheng B. The Association between Red Blood Cell Distribution Width and Mortality in Critically Ill Patients with Acute Kidney Injury. Biomed Res Int. 2018;2018:9658216. Epub 2018/10/23. doi: 10.1155/2018/9658216. PubMed PMID: 30345313; PubMed Central PMCID: PMCPMC6174796.

19. Zou Z, Zhuang Y, Liu L, Shen B, Xu J, Jiang W, et al. Role of elevated red cell distribution width on acute kidney injury patients after cardiac surgery. BMC Cardiovasc Disord. 2018;18(1):166. Epub 2018/08/16. doi: 10.1186/s12872-018-0903-4. PubMed PMID: 30107786; PubMed Central PMCID: PMCPMC6092813.

20. Hu Y, Liu H, Fu S, Wan J, Li X. Red Blood Cell Distribution Width is an Independent Predictor of AKI and Mortality in Patients in the Coronary Care Unit. Kidney Blood Press Res. 2017;42(6):1193–204. Epub 2017/12/12. doi: 10.1159/000485866. PubMed PMID: 29227977.

21. Van Driest SL, Jooste EH, Shi Y, Choi L, Darghosian L, Hill KD, et al. Association Between Early Postoperative Acetaminophen Exposure and Acute Kidney Injury in Pediatric Patients Undergoing Cardiac Surgery. JAMA Pediatr. 2018;172(7):655–63. Epub 2018/05/26. doi: 10.1001/jamapediatrics.2018.0614. PubMed PMID: 29799947; PubMed Central PMCID: PMCPMC6110290.

22. Higgins JP, Thompson SG. Quantifying heterogeneity in a meta-analysis. Stat Med. 2002;21(11):1539–58. doi: 10.1002/sim.1186. PubMed PMID: 12111919.

23. DerSimonian R, Laird N. Meta-analysis in clinical trials. Control Clin Trials. 1986;7(3):177–88. Epub 1986/09/01. PubMed PMID: 3802833.

24. Begg CB, Mazumdar M. Operating characteristics of a rank correlation test for publication bias. Biometrics. 1994;50(4):1088–101. Epub 1994/12/01. PubMed PMID: 7786990.

25. Egger M, Smith GD, Schneider M, Minder C. Bias in meta-analysis detected by a simple, graphical test. BMJ (Clinical research ed). 1997;315(7109):629–34.

26. Akin F, Celik O, Altun I, Ayca B, Ozturk D, Satilmis S, et al. Relation of red cell distribution width to contrast-induced acute kidney injury in patients undergoing a primary percutaneous coronary intervention. Coron Artery Dis. 2015;26(4):289–95. Epub 2015/02/26. doi: 10.1097/mca.0000000000000223. PubMed PMID: 25714066.

27. Elhosseiny S, Akel T, Mroue J, Tathineni P, El Sayegh S, Lafferty J. The Value of Adding Red Cell Distribution Width to Mehran Risk Score to Predict Contrast-induced Acute Kidney Injury in Patients with Acute Coronary Syndrome. Cureus. 2018;10(7):e2911. Epub 2018/09/07. doi: 10.7759/cureus.2911. PubMed PMID: 30186716; PubMed Central PMCID: PMCPMC6122680.

28. Lee SW, Yu MY, Lee H, Ahn SY, Kim S, Chin HJ, et al. Risk Factors for Acute Kidney Injury and In-Hospital Mortality in Patients Receiving Extracorporeal Membrane Oxygenation. PLoS One. 2015;10(10):e0140674. Epub 2015/10/16. doi: 10.1371/journal.pone.0140674. PubMed PMID: 26469793; PubMed Central PMCID: PMCPMC4607159.

29. Mizuno A, Ohde S, Nishizaki Y, Komatsu Y, Niwa K. Additional value of the red blood cell distribution width to the Mehran risk score for predicting contrast-induced acute kidney injury in patients with ST-elevation acute myocardial infarction. J Cardiol. 2015;66(1):41–5. Epub 2014/12/03. doi: 10.1016/j.jjcc.2014.09.006. PubMed PMID: 25448729.

30. Xia Y, Chen D, Li L. Red blood cell distribution width predicts acute kidney injury after cardiac surgery in elderly patients: A retrospective cohort study. Biomedical Research (India). 2017;28(5):1944–9.

31. Zhang L, Guo KP, Mo Y, Yi SW, Huang CZ, Long CX, et al. Predictive value of red blood cell distribution width for acute kidney injury in children with sepsis. Chinese Journal of Contemporary Pediatrics. 2018;20(7):559–62. doi: http://dx.doi.org/10.7499/j.issn.1008-8830.2018.07.009.

32. Wang Y, Bellomo R. Cardiac surgery-associated acute kidney injury: risk factors, pathophysiology and treatment. Nature reviews Nephrology. 2017;13(11):697–711. Epub 2017/09/05. doi: 10.1038/nrneph.2017.119. PubMed PMID: 28869251.

33. O’Connor ME, Kirwan CJ, Pearse RM, Prowle JR. Incidence and associations of acute kidney injury after major abdominal surgery. 2016;42(4):521–30. doi: 10.1007/s00134-015-4157-7. PubMed PMID: 26602784.

34. Mehran R, Dangas GD, Weisbord SD. Contrast-Associated Acute Kidney Injury. New Engl J Med. 2019;380(22):2146–55. doi: 10.1056/NEJMra1805256.

35. Grams ME, Sang Y, Coresh J, Ballew S, Matsushita K, Molnar MZ, et al. Acute Kidney Injury After Major Surgery: A Retrospective Analysis of Veterans Health Administration Data. American journal of kidney diseases: the official journal of the National Kidney Foundation. 2016;67(6):872–80. Epub 2015/09/05. doi: 10.1053/j.ajkd.2015.07.022. PubMed PMID: 26337133; PubMed Central PMCID: PMCPMC4775458.

36. Grams ME, Sang Y, Coresh J, Ballew SH, Matsushita K, Levey AS, et al. Candidate Surrogate End Points for ESRD after AKI. Journal of the American Society of Nephrology: JASN. 2016;27(9):2851–9. Epub 2016/02/10. doi: 10.1681/asn.2015070829. PubMed PMID: 26857682; PubMed Central PMCID: PMCPMC5004655.

37. Hobson CE, Yavas S, Segal MS, Schold JD, Tribble CG, Layon AJ, et al. Acute kidney injury is associated with increased long-term mortality after cardiothoracic surgery. Circulation. 2009;119(18):2444–53. Epub 2009/04/29. doi: 10.1161/circulationaha.108.800011. PubMed PMID: 19398670.

38. Li N, Zhou H, Tang Q. Red Blood Cell Distribution Width: A Novel Predictive Indicator for Cardiovascular and Cerebrovascular Diseases. 2017;2017:7089493. doi: 10.1155/2017/7089493. PubMed PMID: 29038615.

39. Shao Q, Li L, Li G, Liu T. Prognostic value of red blood cell distribution width in heart failure patients: a metaanalysis. Int J Cardiol. 2015;179:495–9. Epub 2014/12/04. doi: 10.1016/j.ijcard.2014.11.042. PubMed PMID: 25465815.

40. Zhang T, Li J, Lin Y, Yang H, Cao S. Association Between Red Blood Cell Distribution Width and All-cause Mortality in Chronic Kidney Disease Patients: A Systematic Review and Meta-analysis. Archives of medical research. 2017;48(4):378–85. Epub 2017/09/17. doi: 10.1016/j.arcmed.2017.06.009. PubMed PMID: 28916240.

41. Xu WY, Yang XB, Wang WQ, Bai Y, Long JY, Lin JZ, et al. Prognostic impact of the red cell distribution width in esophageal cancer patients: A systematic review and meta-analysis. World journal of gastroenterology. 2018;24(19):2120–9. Epub 2018/05/23. doi: 10.3748/wjg.v24.i19.2120. PubMed PMID: 29785080; PubMed Central PMCID: PMCPmc5960817.

42. Tham T, Bardash Y, Teegala S, Herman WS, Costantino PD. The red cell distribution width as a prognostic indicator in upper aerodigestive tract (UADT) cancer: A systematic review and meta-analysis. American journal of otolaryngology. 2018;39(4):453–8. Epub 2018/04/28. doi: 10.1016/j.amjoto.2018.04.013. PubMed PMID: 29699714.

43. Yao D, Wang Z, Cai H, Li Y, Li B. Relationship between red cell distribution width and prognosis in patients with breast cancer after operation: a retrospective cohort study. 2019;39(7). doi: 10.1042/bsr20190740. PubMed PMID: 31262969.

44. Zou Z, Zhuang Y, Liu L, Shen B, Xu J, Jiang W, et al. Role of elevated red cell distribution width on acute kidney injury patients after cardiac surgery. 2018;18(1):166. doi: 10.1186/s12872-018-0903-4. PubMed PMID: 30107786.

45. Wang B, Lu H, Gong Y, Ying B, Cheng B. The Association between Red Blood Cell Distribution Width and Mortality in Critically Ill Patients with Acute Kidney Injury. BioMed Research International. 2018;2018:9658216. doi: http://dx.doi.org/10.1155/2018/9658216.

46. Shema-Didi L, Ore L, Geron R, Kristal B. Is anemia at hospital admission associated with in-hospital acute kidney injury occurrence? Nephron Clinical practice. 2010;115(2):c168–76. Epub 2010/04/22. doi: 10.1159/000312881. PubMed PMID: 20407277.

47. Semba RD, Patel KV, Ferrucci L, Sun K, Roy CN, Guralnik JM, et al. Serum antioxidants and inflammation predict red cell distribution width in older women: the Women’s Health and Aging Study I. Clin Nutr. 2010;29(5):600–4. Epub 2010/03/26. doi: 10.1016/j.clnu.2010.03.001. PubMed PMID: 20334961; PubMed Central PMCID: PMCPMC3243048.

48. Shimazui T, Nakada TA, Tateishi Y, Oshima T, Aizimu T, Oda S. Association between serum levels of interleukin-6 on ICU admission and subsequent outcomes in critically ill patients with acute kidney injury. BMC nephrology. 2019;20(1):74. Epub 2019/03/03. doi: 10.1186/s12882-019-1265-6. PubMed PMID: 30823904; PubMed Central PMCID: PMCPMC6397495.

49. Lippi G, Targher G, Montagnana M, Salvagno GL, Zoppini G, Guidi GC. Relationship between red blood cell distribution width and kidney function tests in a large cohort of unselected outpatients. Scandinavian journal of clinical and laboratory investigation. 2008;68(8):745–8. Epub 2008/07/12. doi: 10.1080/00365510802213550. PubMed PMID: 18618369.

